# Neutrophil-targeted delivery of circSCMH1 unlocks an acute anti-thromboinflammatory function to restore microvascular perfusion in stroke

**DOI:** 10.64898/2026.03.11.711034

**Authors:** Lian Xu, Xinxin Huang, Zhongqiu Zhou, Yanpeng Jia, Shuo Leng, Ying Bai, Bing Han, Honghong Yao

**Author notes:** Correspondence: Honghong Yao. Lian Xu and Xinxin Huang contributed equally to this work.

## Abstract

**Objectives:** Futile recanalization (FR) after endovascular thrombectomy (EVT) is driven by neutrophil-mediated thromboinflammation, yet conventional neuroprotectants fail to address the spatiotemporal complexity of stroke injury. This study aimed to identify the clinical link between circSCMH1, traditionally viewed as a late-stage repair molecule, and FR, while evaluating its hyperacute anti-inflammatory potential via a neutrophil-targeted delivery system.

**Methods:** We analyzed thrombi and plasma from LVO-AIS patients stratified by functional outcomes. We engineered neutrophil-targeted lipid nanoparticles (circSCMH1@pepLNP) for delivery in a mouse transient middle cerebral artery occlusion model. Therapeutic outcomes were evaluated through infarct volume measurement, histological assessment of NETosis, and intravital two-photon microscopy to monitor real-time neutrophil dynamics and capillary stalling.

**Results:** Clinical analyses showed circSCMH1 was significantly downregulated in FR patients, with its levels negatively correlated with NET burden. Furthermore, circSCMH1-negative neutrophils exhibited a higher propensity for NETosis within the thrombus microenvironment. In mice, circSCMH1@pepLNP achieved specific neutrophil delivery, significantly reduced infarct volume, and suppressed the expression of citrullinated histone H3 and neutrophil elastase. Intravital two-photon and laser speckle imaging further confirmed that this intervention attenuated neutrophil adhesion, increased migratory velocity, and alleviated capillary stalling, thereby restoring cortical microvascular perfusion.

**Conclusions:** Our findings associate neutrophil-specific circSCMH1 downregulation with clinical FR in LVO-AIS. Targeted intracellular delivery reveals an acute anti-thromboinflammatory function that complements established reparative properties, proposing a novel adjunctive approach for EVT to address the spatiotemporal complexity of stroke and improve clinical outcomes.

## Introduction

Acute ischemic stroke (AIS), particularly complicated by large vessel occlusion (LVO), represents a major cause of morbidity and mortality.^1^ Although endovascular thrombectomy (EVT) achieves high macrovascular recanalization, a substantial proportion of patients fail to attain favorable functional outcomes, a clinical phenomenon termed futile recanalization (FR).^2,3^ The pathogenesis of FR is critically driven by post-reperfusion thromboinflammation.^4^ During the hyperacute phase, early infiltrating neutrophils undergo aberrant activation and release neutrophil extracellular traps (NET). This process directly precipitates microvascular thrombosis and capillary stalling, severely impeding tissue-level reperfusion and exacerbating the ischemic core despite the patency of major cerebral arteries.^5–7^

Over the past decades, numerous neuroprotective agents have demonstrated efficacy in preclinical models but failed in clinical translation. A primary limitation of most previously tested systemic pharmacological neuroprotectants is their inability to address the spatiotemporal complexity of ischemic injury.^8–10^ The progression of ischemic stroke is inherently biphasic, characterized by an acute phase dominated by robust thromboinflammatory damage and a delayed phase requiring sustained neurovascular repair.^11^ Consequently, therapies targeting a single pathophysiological event within a narrow time window often yield suboptimal global efficacy. Developing an adjunctive therapy for EVT that can simultaneously mitigate early microvascular thromboinflammation and promote long-term tissue recovery remains a critical unmet clinical need.

Circular RNAs (circRNAs) are highly stable, endogenous transcripts that have emerged as pivotal regulators in the central nervous system.^12^ Among these, circSCMH1 has been extensively documented as a promising therapeutic agent for post-stroke recovery. Existing studies highlight its efficacy in promoting late-stage neural repair, enhancing neuroplasticity, and stimulating angiogenesis.^13–15^ However, its potential application during the hyperacute reperfusion phase, specifically its capacity to modulate innate immune responses and early thromboinflammation, remains completely unexplored. We hypothesized that a fundamental delivery constraint prevents systemic circSCMH1 from reaching its crucial cellular targets in the hyperacute phase, namely, activated neutrophils, the primary effectors of early injury.

To overcome this limitation, we engineered a targeted cellular delivery system utilizing formyl peptide receptor (FPR)-targeting lipid nanoparticles (pepLNP) to specifically direct circSCMH1 into ischemic neutrophils. In this study, we identified that FR is intrinsically associated with the depletion of circSCMH1 within thrombus-associated neutrophils. Crucially, the targeted early delivery of circSCMH1@pepLNP unexpectedly conferred a potent, previously unrecognized anti-thromboinflammatory function upon this transcript. This targeted intervention efficiently abolished NETosis, alleviated microvascular obstruction, and restored cerebral perfusion in stroke models. Given the well-documented role of circSCMH1 in late-stage neurovascular repair, this study unlocks a previously unrecognized hyperacute anti-thromboinflammatory function. Collectively, this targeted early intervention strategy paves the way for a single-formulation, dual-phase therapeutic paradigm for the comprehensive management of acute ischemic stroke.

## Methods

### Human subjects

Patients with large-vessel occlusion acute ischemic stroke (LVO-AIS) were enrolled from Zhongda Hospital, School of Medicine, Southeast University, Nanjing, China. This study was approved by the Institutional Review Board (IRB) and Ethics Committee of Zhongda Hospital, School of Medicine, Southeast University, Nanjing, China (approval ID: 2025ZDSYLL046). The inclusion criteria for this study were as follows: 1) occlusion of the M1/M2 segment of the middle cerebral artery; 2) age > 50 years; 3) undergoing mechanical thrombectomy. Patients with a history of cerebral hemorrhage, malignancy, hematological diseases, inflammatory diseases, or infectious disorders were excluded from the study. Irreversibly infarcted tissues (infarct volume) were defined as the region with relative cerebral blood flow (CBF) < 30% of that in the contralateral normal brain tissue. Hypoperfusion volume was defined as the tissue with a Tmax value > 6 s. Automatic volumes of the ischemic core and hypoperfusion region were calculated using these thresholds. The mismatch volume (penumbra volume) was computed as the hypoperfusion volume minus the ischemic core volume. Good functional outcome at 3 months was defined as a modified Rankin Scale (mRS) score of 0–2, whereas poor functional outcome was defined as an mRS score of 3–6.

In this study, the thrombus and plasma samples were collected from LVO-AIS patients subjected to mechanical thrombectomy. These patients were classified into MR and FR groups after a 3-month follow-up. All subjects provided informed consent for sample collection.

### Immunostaining of human thrombus

Retrieved thrombi were promptly fixed in 10% formalin and subsequently embedded in paraffin. These formalin-fixed paraffin-embedded samples were cut into 5 μm sections, after which deparaffinization was performed via successive immersion in xylene I (15 minutes) and xylene II (15 minutes). Subsequently, graded ethanol rehydration was conducted in the following order: 100% ethanol I (5 minutes), 100% ethanol II (5 minutes), 90% ethanol (5 minutes), 80% ethanol (5 minutes), 70% ethanol (5 minutes), and 50% ethanol (5 minutes), with the process concluding with two 5-minute rinses in distilled water.

Antigen retrieval was carried out by incubating the sections in preheated (95–100[) 1 × Sodium citrate antigen retrieval solution (P0083, Beyotime) in a water bath for 20 minutes, followed by slow cooling to room temperature (20–30 minutes). The sections were blocked for 2 hours at room temperature with PBS supplemented with 10% normal goat serum (NGS) and 0.25% Triton X-100 (1139ML100, Biofroxx). Primary antibodies, namely anti-Neutrophil Elastase (NE) (1:200, TA0010s, Abmart) and anti-H3Cit (1:200, AB5103, Abcam), were added to the blocking buffer and incubated overnight at 4[. Following three 5-minute rinses with PBS, the sections were incubated with species-compatible fluorescent secondary antibodies (Alexa Fluor 488 goat anti-rabbit IgG, 1:200, A-11008, Invitrogen) for 1 hour at room temperature. After three additional 5-minute PBS rinses, the sections were coverslipped using DAPI-containing antifade mounting medium (0100-20, SouthernBiotech). The prepared slides were stored at 4[in a dark environment to preserve fluorescence prior to imaging. High-resolution z-stack imaging (1 μm optical sections) was performed using a whole-slide scanner (DML8, Leica) and laser scanning confocal microscopes (Leica STELLARIS, Leica).

### Enzyme-linked immunosorbent assay (ELISA)

Blood samples were collected from LVO-AIS patients before they underwent thrombectomy. All plasma samples were isolated in the clinical laboratories of the participating hospitals and promptly stored at −80°C until laboratory detection. The concentrations of plasma H3cit and MPO-DNA were determined using commercially available ELISA kits [Human H3cit (501620, Cayman) and Human MPO-DNA (YJ342558A, YUANJU BIO)] in strict accordance with the manufacturer’s protocols, consistent with previous descriptions.^16^

### Animals

Unless stated otherwise, C57BL/6J mice, aged 8 to 10 weeks were purchased from GemPharmatech (Nanjing, China). All animals were housed under standardized environmental conditions, which included regulated temperature and relative humidity, a 12-hour light/12-hour dark cycle (with illumination starting at 7:00 AM), and ad libitum access to standard chow and drinking water. The experimental protocol was reviewed and approved by the Institutional Animal Care and Use Committee (IACUC) of the Medical School, Southeast University (Nanjing, Jiangsu Province, China; approval identification number: 20260101201). All in vivo experiments were conducted in strict compliance with the ARRIVE (Animal Research: Reporting of In Vivo Experiments) guidelines.

### Transient middle cerebral artery occlusion (tMCAO)

Focal cerebral ischemia was established via the intraluminal filament method in accordance with well-established experimental procedures. Mice were anesthetized by inhalation of 3% isoflurane for induction, and anesthesia was maintained with 1.5% isoflurane mixed with 30% oxygen and 70% nitrous oxide, which was delivered through a face mask. Following dissection and exposure of the right external carotid artery (ECA), a silicone-coated 6-0 nylon monofilament (Doccol, USA) was inserted into the lumen of the ECA and advanced approximately 9–10 mm along the internal carotid artery to occlude the origin of the middle cerebral artery (MCA). Cerebral blood flow was reestablished after 60 minutes of occlusion by withdrawing the filament. Throughout the entire surgical process, a feedback-controlled heating system was used to maintain the core body temperature of the mice at 37.0 ± 0.5[. Infarct volume was assessed at 24 hours after reperfusion.

### Nissl staining and brain atrophy measurement

Coronal brain sections with a thickness of 30 μm were fabricated using a cryostat microtome. Every sixth section was harvested throughout the entire infarct area. Histological evaluation of infarct volume was performed using Nissl staining (G1036-100 mL, Servicebio), strictly following the manufacturer’s instructions. Systematic sampling was implemented by examining every sixth section that spanned the whole infarct region. Digital images of the stained sections were captured under standardized illumination conditions. Atrophy volume was calculated as the difference between the contralateral brain area and the ipsilateral brain area. Bright-field images were acquired at 10× magnification using a whole-slide scanner (DML8, Leica) and a digital section scanning microscope (VS200, Olympus). All data analyses were conducted with Image J software.

### Immunostaining and image analysis of brain sections

Mice were transcardially perfused with PBS, followed by pre-fixation in 4% paraformaldehyde (PFA). Brain tissues were then harvested and subjected to further fixation for 24 hours. Subsequently, the tissues were transferred into 30% sucrose solution and incubated until they sank to the bottom of the solution. Coronal brain sections (30 μm thick) were prepared with a cryostat and rinsed three times with PBS (5 minutes per rinse).

The sections were then blocked with 5% bovine serum albumin (BSA) dissolved in PBST for 1 hour at room temperature. Primary antibodies were diluted in PBST and incubated with the sections overnight at 4[under gentle agitation. The primary antibodies employed were as follows: anti-NE (1:200, TA0010s, Abmart), anti-CD31 (1:50, AF3628, RD Systems), anti-H3Cit (1:200, AB5103, Abcam), anti-F7/4 (1:50, AB53453, Abcam), and anti-Fibrin (1:200, AB27913, Abcam). After primary antibody incubation, the sections were washed three times with PBST (10 minutes per wash). Corresponding Alexa Fluor-conjugated secondary antibodies (all diluted to 1:200 in PBST) were added and incubated for 2 hours at room temperature with gentle shaking. The secondary antibodies used included: Alexa Fluor 647 donkey anti-rabbit IgG (1:200, A-31573, Invitrogen), Alexa Fluor 594 donkey anti-Goat IgG (1:200, A-11058, Invitrogen), and Alexa Fluor 488 goat anti-Rat IgG (1:200, A-48262, Invitrogen). Excess staining and unbound antibodies were removed by three additional PBST washes (10 minutes per wash). Finally, the sections were mounted on glass slides using an antifade mounting medium containing DAPI (0100-20, Southern Biotech) for imaging. Images were acquired using a confocal microscope (Leica STELLARIS, Leica) and analyzed with ImageJ (1.54f) and Imaris (9.0.1) software.

### Western blot analysis

Brain tissues and cells were homogenized thoroughly in RIPA lysis buffer (P0013B, Beyotime) supplemented with a cocktail of protease inhibitors. After quantification of protein lysates for concentration, equal quantities of proteins were separated by SDS-PAGE and then transferred electrophoretically onto polyvinylidene fluoride (PVDF) membranes. The membranes were blocked with 5% non-fat milk dissolved in TBST for 1 hour at room temperature, followed by overnight incubation at 4 °C with primary antibodies specific to the following targets: NE (1:1000, TA0010s, Abmart), H3cit (1:1000, AB5103, Abcam), Fibrinogen (1:1000, AB27913, Abcam), CD41 (1:1000, GTX113758, GeneTex), and β-actin (1:1000, 66009-1-Ig, Proteintech). Following three washes with TBST, the membranes were incubated for 1 hour with horseradish peroxidase (HRP)-conjugated goat anti-mouse/rabbit secondary antibodies (1:2000, 7076P2/7074P2, Cell Signaling Technology). After three additional TBST washes, protein bands were detected using an automated chemiluminescence detection system (Tanon 5200, Tanon Science & Technology), and densitometric quantification was performed using ImageJ (1.54f) software. Protein expression levels were normalized against β-actin, which served as internal loading controls.

### Preparations of circSCMH1@pepLNP

All LNP formulations were fabricated via a standard ethanol dilution approach. Specifically, ionizable lipid, helper lipid DSPC, cholesterol, DSPE-PEG, and DSPE-PEG-pep were dissolved into ethanol at a molar ratio of 50:38.5:10:0.5. Mouse circscmh1 was synthesized by GenePharma Co., Ltd. (Shanghai, China). We prepared the aqueous phase by dissolving circSCMH1 in 10 mM citrate buffer with a pH of 4.5, following the corresponding protocol. Next, the ethanol phase containing the lipid mixture and the aqueous phase with circSCMH1 were vortexed at a speed of 1,500 rpm for 1 minute at a volume ratio of 1:3, and then incubated at room temperature for about 30 minutes to form circSCMH1@pepLNP. After that, the obtained circSCMH1@pepLNP dispersion was diluted in PBS and subjected to ultrafiltration to eliminate residual ethanol. The resulting circSCMH1@pepLNP was filtered through a 0.22-μm filter for sterilization. For preparing the control group (circSCMH1@LNP), we adopted the identical procedure, with the only difference being that DSPE-PEG-pep was excluded from the lipid phase.

### Intravital imaging of the microvasculature and neutrophil dynamics

#### Cranial window implantation

Cranial window implantation surgery was performed on 8–10-week-old C57BL/6 mice as previously reported[16]. In brief, a circular craniotomy with a diameter of 5 mm was drilled over the right somatosensory cortex, with the center coordinates set at AP: -1.80 mm and ML: -1.80 mm relative to bregma. A sterile glass coverslip was then inserted into the craniotomy site and affixed using tissue adhesive. At the same time, a head-plate was attached to the skull area surrounding the craniotomy and immobilized with dental cement. Following the completion of surgery, the mice were housed for a two-week recovery period before being subjected to subsequent experimental procedures.

#### Laser speckle cortical imaging

Laser speckle contrast imaging (LSCI) was performed on awake mice. First, we immobilized the mice using a customized body holder. To avoid ocular dryness during imaging, erythromycin eye ointment was gently applied to their eyes. The mice were then precisely positioned under the LSCI system (RWD, RFLSI Ⅲ) and given a 5-minute acclimatization period to adapt to the environment. After acclimatization, relative cerebral blood flow (CBF) changes within the cranial window were quantitatively analyzed using LSCI software (V01.00.05.18305, RWD). We set the imaging parameters as follows: a resolution of 3.4 µm/pixel, a frame rate of 2 Hz, and a laser intensity of 110 mW. The region of interest (ROI) was selected within the territory of the terminal branches of the middle cerebral artery (MCA). Specifically, we defined the ROI as a circle (200 µm in diameter) with the middle cerebral artery passing through its center. The perfusion signal detected within this ROI was used to reflect the blood perfusion status of the MCA territory.

#### Two-photon imaging of neutrophils in microvessels

Two-photon imaging (TPM) was carried out using a TPM system integrated with a femtosecond fiber laser (with approximately 35 mW power at the objective; TVS-FL-01, Transcend Vivoscope Biotech Co., Ltd). During the imaging process, the mice were head-fixed in an adapter and remained conscious throughout. To visualize the cerebral vasculature, we intravenously injected FITC-dextran (5% w/v, 50 µL; FD70S, Sigma Aldrich) into the tail vein. Neutrophils were labeled with Rhodamine 6G (1 mg/mL, 100 µL; 83697, Sigma Aldrich). Both dyes were administered intravenously 15 minutes prior to imaging to ensure adequate circulation and effective labeling. The gap between the objective lens and the glass coverslip was filled with 1.5% low-melting-point agarose. Image acquisition and microscope control were managed using GINKGO-MTPM software (version 1.0.28, Transcend Vivoscope Biotech Co., Ltd). ROIs in the penumbra region were selected for TPM imaging based on the blood flow changes detected by LSCI. All images were acquired at a frame rate of 10 Hz, with a resolution of 512 × 512 pixels and a field of view of 420 µm × 420 µm. Z-stacks (x,y,z images) were recorded in the selected regions with a step size of 1 µm, a resolution of 400 × 400 pixels, and a frame rate of 4 Hz, covering an approximate volume of 230 × 230 × 200 µm³. Capillary stalls were defined as neutrophils remaining stationary for more than 5 seconds at sites not stained by FITC-dextran. After data collection, images were processed and analyzed using Fiji and MATLAB software (MathWorks). Mean track velocities (µm/s) and track durations (s) were calculated for individual tracks lasting 5 minutes. All track calculations were performed using the TrackMate plugin in Fiji.

## Results

### circSCMH1 is decreased in LVO-AIS patients with FR and negatively correlates with NET burden

To investigate the clinical relevance of circSCMH1 in post-stroke thromboinflammation, we prospectively enrolled a cohort of LVO-AIS patients who underwent endovascular thrombectomy. Thrombus and peripheral blood samples were collected, and patients were subsequently dichotomized into meaningful recanalization (MR) and FR groups based on their 3-month functional outcomes (**Figure 1A**). Quantitative PCR (qPCR) analysis revealed that circSCMH1 expression was significantly downregulated in both the thrombi and plasma of the FR group compared to the MR group (**Figure 1B-C**). Notably, plasma circSCMH1 levels exhibited a significant negative correlation with circulating NET biomarkers (**Figure 1D**).

**Figure 1.**
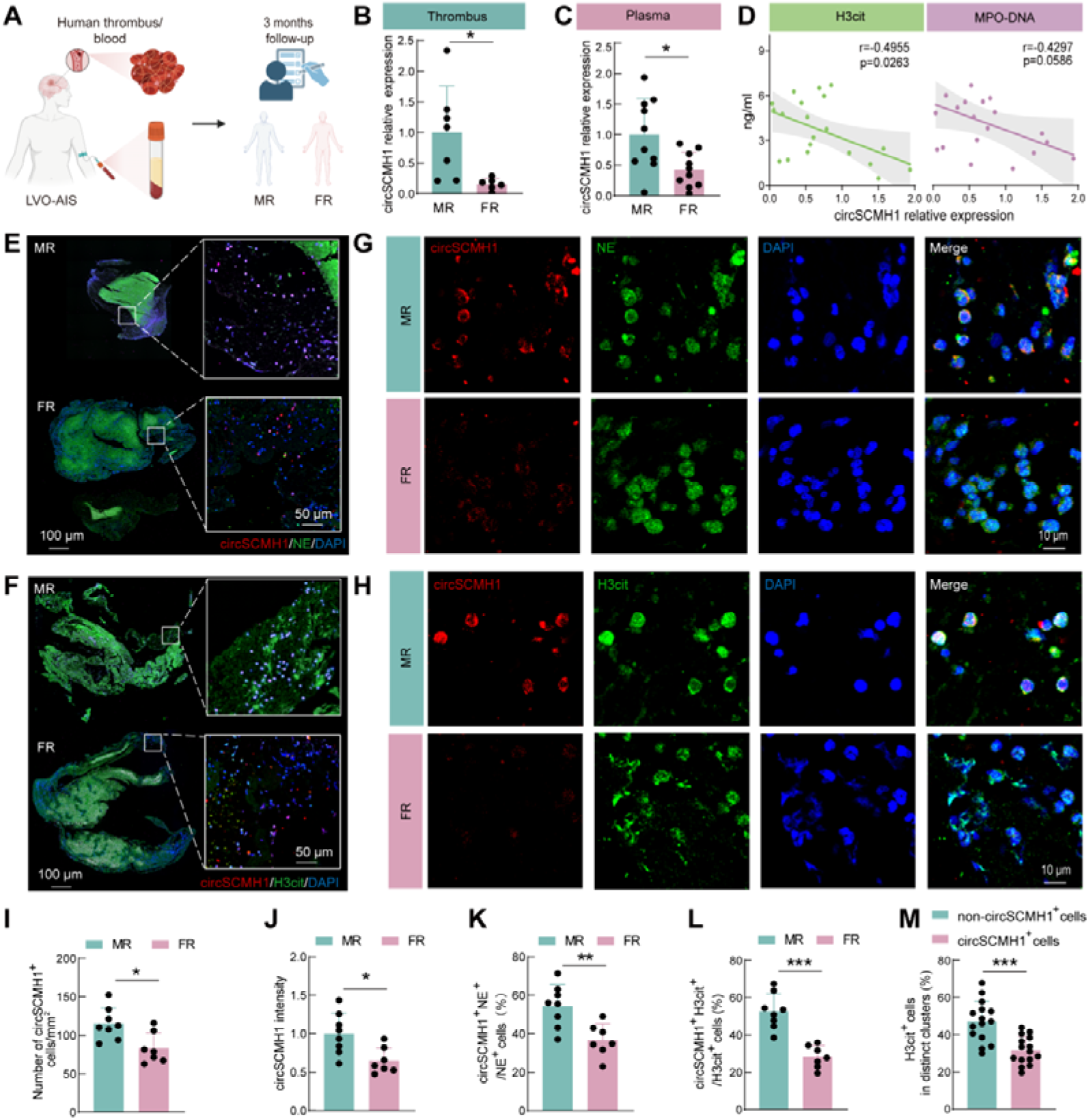
circSCMH1 is significantly decreased in LVO-AIS patients with FR and negatively correlates with NETs burden. (**A**) Schematic of the experimental design for clinical sample collection. Thrombus and plasma were collected from LVO-AIS patients. The patients were divided into meaningful recanalization (MR) and futile recanalization (FR) groups based on their 3-month functional outcomes. (**B-C**) qPCR analysis of circSCMH1 expression in Thrombus (B) and plasma (C) samples obtained from LVO-AIS patients before thrombectomy. MR group: thrombus, n=7; plasma, n=10. FR group: thrombus, n=6; plasma, n=10. \**p*<0.05 using Student’s *t* test. (**D**) Correlation analysis between relative plasma circSCMH1 expression and circulating levels of NET biomarkers, H3Cit (left) and MPO-DNA complexes (right) in (C). (**E-F**) Representative low-magnification immunofluorescence images of thrombus sections from MR and FR patients stained for circSCMH1 (red), NE (green, E) and H3cit (green, F). (**G-H**) Representative high-resolution immunofluorescence images of thrombus sections from MR and FR for circSCMH1 (red), NE (green, G) and H3cit (green, H). (**I**) Quantification of the average count of circSCMH1^+^ cells (red) in thrombus from LVO-AIS patients. MR group: n=8; FR group: n=7. \**p*<0.05 using Student’s t test. (**J**) Quantitation of circSCMH1 relative fluorescence intensity. MR group: n=8; FR group: n=7. \**p*<0.05 using Student’s t test. (**K**) Quantitative analysis of the percentage of circSCMH1^+^ neutrophils in the thrombus of MR and FR patients. MR group: n=8; FR group: n=7. \*\**p*<0.01 using Student’s t test. (**L**) Quantitative analysis of the percentage of circSCMH1^+^H3Cit^+^ cells in the thrombus of MR and FR patients. MR group: n=8; FR group: n=7. \*\*\**p*<0.001 using Student’s t test. (**M**) The proportions of H3Cit^+^ cells were compared in circSCMH1^+^ neutrophils versus in non-circSCMH1^+^ neutrophils. n=15 individuals per group. \*\*\**p*<0.001 using Student’s t test. LVO-AIS, large-vessel occlusion acute ischemic stroke. MR, meaningful recanalization. FR, futile recanalization. MPO-DNA,DNA complexes with myeloperoxidase. NE, neutrophil elastase.

The involvement of circSCMH1^+^ neutrophils in NET formation and their subsequent association with FR was investigated through the analysis of thrombectomy-retrieved thrombi (**Figure 1E-H**). The circSCMH1-positive cells observed in the thrombi of MR patients were significantly reduced in the thrombi of FR patients and the FR group showed lower fluorescence intensity of circSCMH1 (**Figure 1I-J**). Congruently, a significantly lower proportion of circSCMH1^+^ neutrophils and circSCMH1^+^ NETs within the FR group (**Figure 1K-L**). Most importantly, at the single-cell level, circSCMH1^+^ neutrophils exhibited a significantly lower propensity to undergo NETosis (indicated by H3Cit positivity) compared to their circSCMH1-negative counterparts within the same thrombus (**Figure 1M**). Collectively, these clinical findings indicate that the downregulation of endogenous circSCMH1 in neutrophils is intrinsically linked to exacerbated NET-driven thromboinflammation and the subsequent development of FR.

### Targeted delivery of circSCMH1 to neutrophils significantly mitigates thromboinflammation post-reperfusion in tMCAO mice

To realize targeted in vivo delivery of circSCMH1 to neutrophils, we engineered a neutrophil-specific lipid nanoparticle (LNP) delivery system loaded with circSCMH1, designated circSCMH1@pepLNP (**Figure 2A**), with the cFLFLF motif for specific targeting of formyl peptide receptors (FPRs) expressed on neutrophils. Transmission electron microscopy (TEM) analysis verified the generation of monodispersed, spherical nanoparticles (**Figure 2B**). Following intravenous administration, immunofluorescence imaging at 24 hours post-injection revealed a markedly enhanced accumulation of circSCMH1 in the peripheral blood neutrophils of mice receiving circSCMH1@pepLNP (**Figure 2C**). Consistent with this, qPCR assays demonstrated a significant elevation of circSCMH1 expression in these isolated neutrophils (**Figure 2D**). Collectively, these results verify the efficient, cell-specific delivery capacity of the engineered pepLNP system.

**Figure 2.**
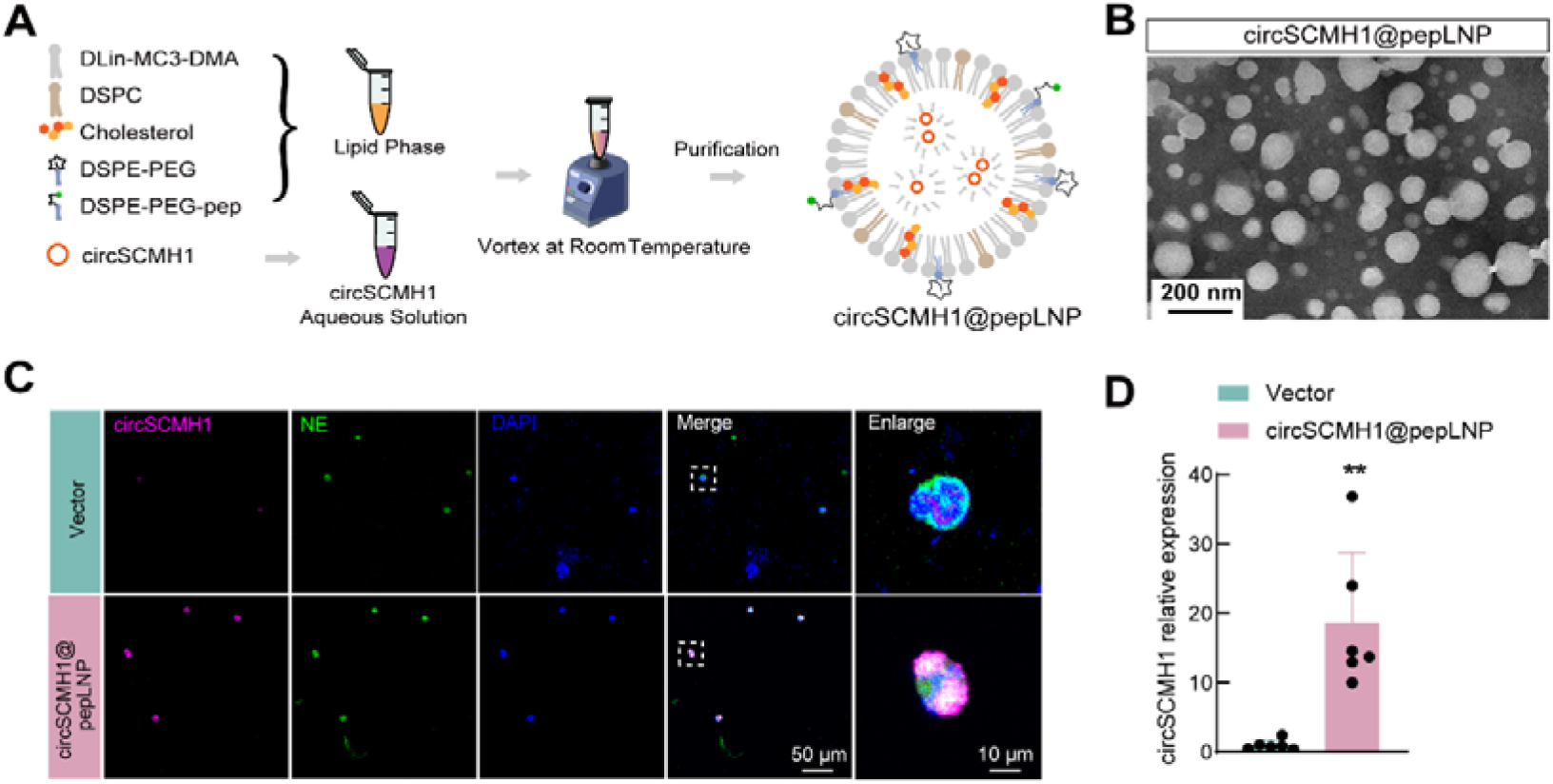
Preparation and neutrophil-targeting capacity of circSCMH1@pepLNP. **(A)** Schematic illustration of the preparation process for circSCMH1@pepLNP. The lipid phase, incorporating the formyl peptide receptor (FPR)-targeting peptide (DSPE-PEG-pep) and the ionizable lipid, is rapidly mixed with the circSCMH1 aqueous solution to assemble the targeted nanoparticles. **(B)** Representative TEM image displaying the spherical morphology and uniform size distribution of the engineered circSCMH1@pepLNP. Scale bar: 200 nm. **(C)** Representative immunofluorescence images of peripheral blood neutrophils isolated from tMCAO mice following intravenous injection of either Vector control or circSCMH1@pepLNP. Cells were stained to detect circSCMH1 (magenta) and the specific neutrophil marker NE (green). Nuclei were counterstained with DAPI (blue). **(D)** Quantitative qPCR analysis of circSCMH1 expression in peripheral blood neutrophils obtained from tMCAO mice. Vector group: n=6; circSCMH1@pepLNP group: n=6. \*\**p*<0.01 using Student’s t test. FPR, formyl peptide receptor; TEM, transmission electron microscopy; NE, neutrophil elastase.

To evaluate the therapeutic potential of early post-reperfusion targeted delivery of circSCMH1 to neutrophils, we assessed the impact of circSCMH1@pepLNP on ischemic brain injury. Nissl staining circSCMH1@pepLNP treatment significantly reduced infarct volume at day 3 post-reperfusion compared to the Vector control. (**Figures 3A**). Moreover, the therapeutic intervention significantly reduced the burden of platelet deposition, as evidenced by decreased protein levels of CD41 and fibrin(ogen) observed at day 3 post-reperfusion (**Figures 3B**). Additionally, the formation of NET was markedly inhibited, confirmed by the reduced expression of neutrophil elastase (NE) and citrullinated histone H3 (H3Cit) (**Figures 3C**). Concurrently, the expression levels of these proteins were consistently decreased at both day 1 and day 3 following tMCAO **(Supplemental Figure 1)**.

**Figure 3.**
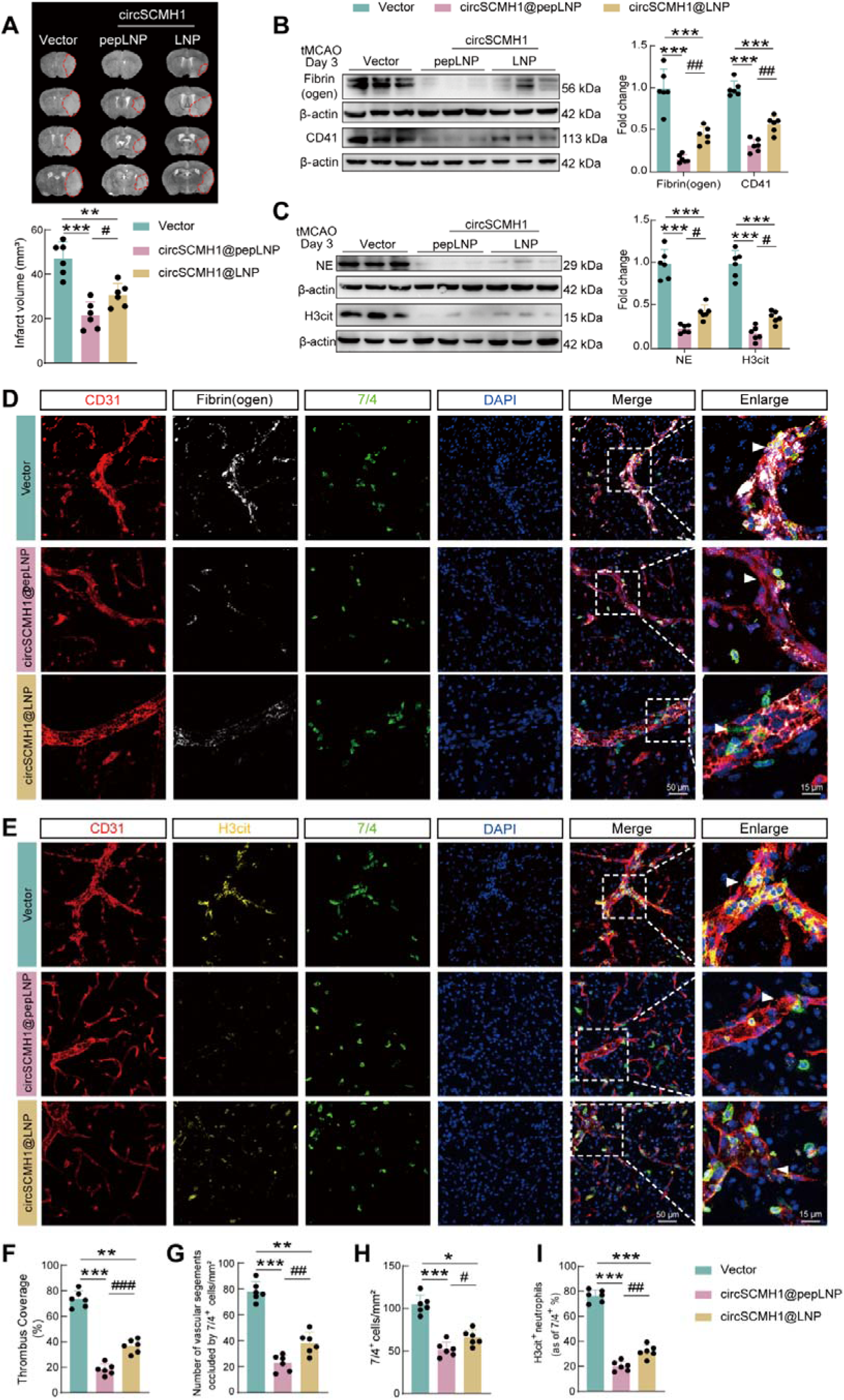
Targeted delivery of circSCMH1 to neutrophils significantly reduced thromboinflammation post-reperfusion in tMCAO mice. (**A**) Representative images of Nissl-stained coronal brain sections and quantification of infarct volume (outlined in red) at day 3 post-reperfusion in mice treated with Vector, circSCMH1@pepLNP, or circSCMH1@LNP. n=6 mice per group. \*\**p*<0.01 and \*\*\**p*<0.001 versus the Vector group, ^#^*p*<0.05 versus the circSCMH1@pepLNP group using one-way ANOVA followed by Sidak post hoc multiple comparisons test. (**B-C**) Representative Western blots and quantitative analysis of thrombosis markers (Fibrin(ogen) and CD41, B) and NETosis markers (NE and H3Cit, C) in ischemic cortex at day 3 post-reperfusion. \*\*\**p*<0.001 versus the Vector group, ^#^*p*<0.05 and ^##^*p*<0.01 versus the circSCMH1@pepLNP group using one-way ANOVA followed by Sidak post hoc multiple comparisons test. (**D**) Representative immunofluorescence images of thrombosed capillaries (CD31^+^ red, fibrinogen^+^ white, 7/4^+^ green) in the ischemic cortex at day 3 post-reperfusion. (**E**) Representative immunofluorescence images of stalled neutrophils and NETs (CD31^+^ red, H3cit yellow, 7/4^+^ green) in the ischemic cortex of tMCAO mice at day 3 post-reperfusion. (**F**) Quantification of thrombus coverage (ratio of fibrinogen-positive area to CD31-positive area). \*\**p*<0.01 and \*\*\**p*<0.001 versus the Vector group, ^###^*p*<0.001 versus the circSCMH1@pepLNP group using one-way ANOVA followed by Sidak post hoc multiple comparisons test. (**G-H**) Quantification of neutrophils (7/4^+^, green) and vascular segments (CD31^+^ red) occluded by neutrophils (7/4^+^ green). \**p*<0.05, \*\**p*<0.01, and \*\*\**p*<0.001 versus the Vector group, ^#^*p*<0.05 and ^##^*p*<0.01 versus the circSCMH1@pepLNP group using one-way ANOVA followed by Sidak post hoc multiple comparisons test. (**I**) Quantification of the proportion of neutrophils that produce extracellular traps (ratio of H3cit^+^7/4^+^ cells to all 7/4^+^ cells). \*\*\**p*<0.001 versus the Vector group, ^##^*p*<0.01 versus the circSCMH1@pepLNP group using one-way ANOVA followed by Sidak post hoc multiple comparisons test. NE, neutrophil elastase. NETs, neutrophil extracellular traps.

Immunofluorescence staining further corroborated these biochemical findings, showing that circSCMH1@pepLNP treatment substantially diminished microthrombus formation, reduced the density of infiltrating neutrophils (7/4+), and decreased the occlusion of cerebral microvessels (**Figures 3E-H**). Furthermore, the proportion of neutrophils undergoing NETosis within the total infiltrating neutrophil population was significantly reduced (**Figures 3I**), indicating an intrinsic suppression of NET formation by the intracellular delivery of circSCMH1. Collectively, our findings suggest that targeted neutrophil delivery of circSCMH1 can significantly reduce the thrombotic inflammatory response after reperfusion, thereby improving the no-reflow phenomenon, and the improvement effect is significantly superior to that of non-targeted delivery.

### CircSCMH1@pepLNP restores cerebral blood flow and attenuates neutrophil adhesion to capillaries

To identify no-reflow microvessels *ex vivo*, we used an *in vivo* labeling approach. Mice were intravenously injected with two distinct fluorescent *Lycopersicon esculentum* (tomato) lectins (LEL) before ischemia and after reperfusion, respectively. Specifically, the first injection (red LEL) was administered 5 minutes prior to the establishment of the tMCAO model to label all cerebral blood vessels, while the second injection (green LEL) was given 5 minutes before sacrifice to label functionally perfused blood vessels. Microvessels that were only labeled by the first tomato lectin (red LEL) but not by the second one (green LEL) were defined as no-reflow microvessels (**Figures 4A**). Representative fluorescence images of 50-μm thick brain slices revealed that in the Vector-treated group, only major vessels achieved recanalization following reperfusion. In contrast, the proportion of no-reflow microvessels was significantly reduced in the circSCMH1@pepLNP group (**Figures 4B-D**). Parallel assessment via quantitative fluorescence imaging demonstrated a substantial reduction in the area fraction of non-reflow vessels within the infarcted cortex of circSCMH1@pepLNP-treated mice **(Supplemental Figure 2)**. Consistent with this these microvascular findings, wide-field cortical perfusion monitoring by laser speckle contrast imaging (LSCI) confirmed macroscopic therapeutic efficacy, showing that circSCMH1@pepLNP significantly improved overall cerebral reperfusion at both day 1 and day 3 post-tMCAO compared to the Vector group (**Figures 4E-F**).

**Figure 4.**
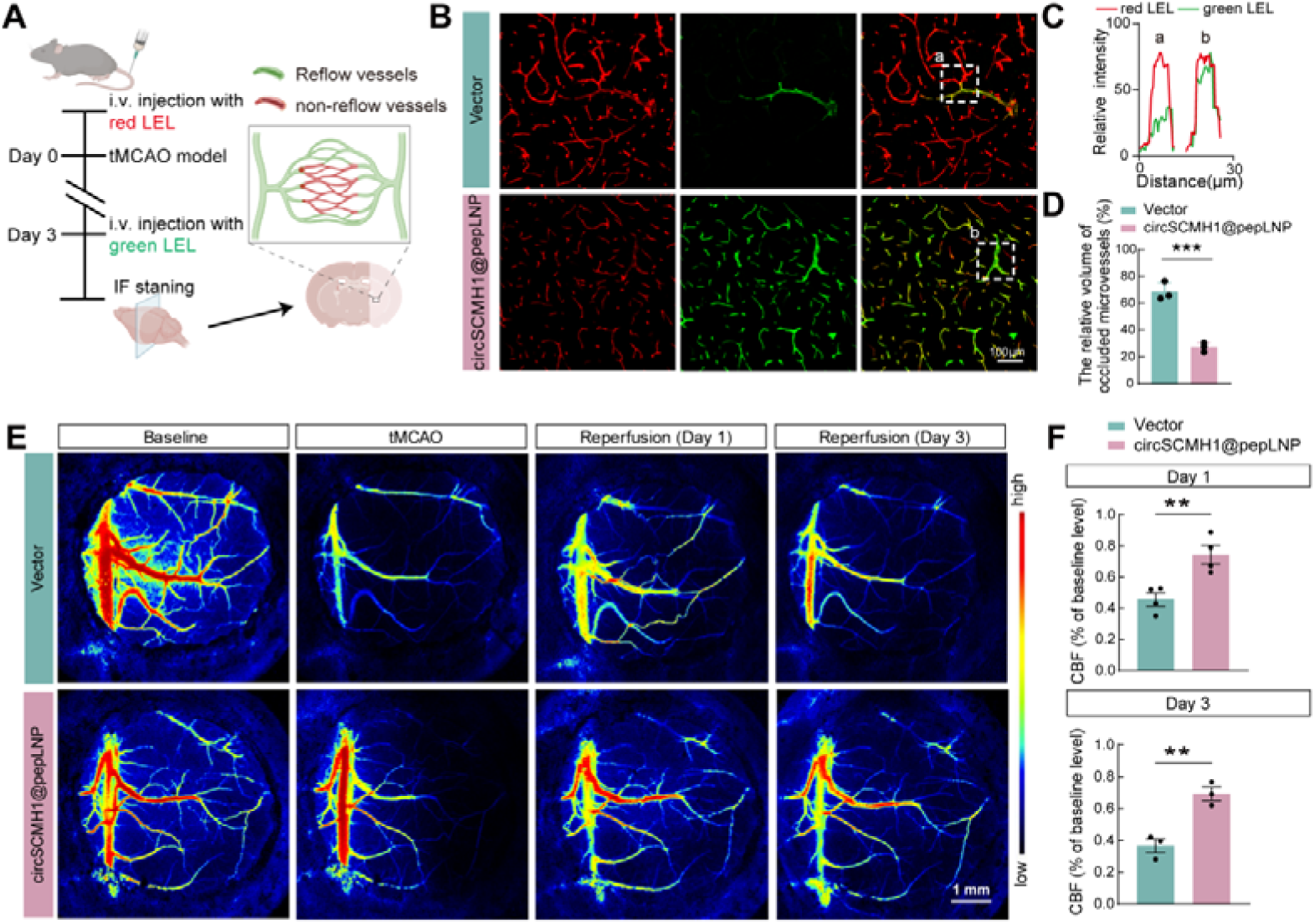
Targeted delivery of circSCMH1@pepLNP restored cerebral blood flow and cortical cerebral blood flow following tMCAO. (**A**) Schematic illustration of the *in vivo* dual-lectin perfusion labeling strategy used to identify the microvascular no-reflow phenomenon. A red-fluorescent *Lycopersicon esculentum* lectin (red LEL) was injected intravenously prior to ischemia to label the entire baseline vascular network, followed by a green LEL injection post-reperfusion to label only the functionally perfused vessels. Consequently, occluded (non-reflow) microvessels are exclusively labeled by the red LEL without overlapping green signal. (**B**) Representative high-resolution fluorescence images of the ischemic cerebral cortex from Vector and circSCMH1@pepLNP-treated mice at day 3 post-tMCAO, distinguishing recanalized (yellow, merged) from non-reflow (red only) microvessels. (**C**) Normalized intensity profiles of the marked white boxes (a and b) marker in (B). The red lines represent the signal from red LEL, and the green lines denote the signal from green LEL. (**D**) Quantification of the relative volume of occluded microvessels after reperfusion. n=6 mice per group. \*\*\**p*<0.001 using Student’s t test. (**E-F**) Representative Laser Speckle Contrast Imaging (LSCI) images illustrating cortical perfusion in tMCAO mice following administration of Vector or circSCMH1@pepLNP (E) and quantitative temporal analysis (F) of macroscopic cortical cerebral blood flow (CBF) at baseline, during MCAO, and at days 1 and 3 post-reperfusion. n=4 mice per group at Day 1 and n=3 mice per group at Day 3. \*\**p*<0.01 using Student’s t test. Scale bar:1 mm. CBF, cerebral blood flow. FOV, field of view. LEL, lycopersicon esculentum (tomato) lectin. LSCI, laser speckle cortical imaging.

Ischemia-induced neutrophil adhesion to the vascular endothelium is a primary driver of capillary obstruction and subsequent microvascular hypoperfusion. To directly visualize the regulatory role of circSCMH1 in neutrophil dynamics under no-reflow conditions, we conducted real-time intravital imaging through a cranial window, which specifically targeted the distal (pial) segments of the middle cerebral artery (MCA) (**Figures 5A**). Utilizing in vivo two-photon microscopy, we labeled neutrophils with Rhodamine6G and vascular structures with Dextran, thereby enabling detailed characterization of neutrophil behaviors. Systemic administration of circSCMH1@pepLNP reperfusion remarkably altered neutrophil dynamics, which was reflected in three key aspects: i) a significant reduction in neutrophil rolling and adhesion (**Figure 5B-C**), ii) a pronounced increase in migratory velocity, and iii) a shortened duration of capillary stalling (**Figure 5D-E**). Quantitative mapping of microvascular patency further confirmed that targeted circSCMH1 delivery effectively alleviated neutrophil-mediated capillary plugging, thereby preserving microcirculatory flow (**Figure 5D-E**). Collectively, these *in vivo* findings establish circSCMH1 as a potent modulator of neutrophil trafficking and microvascular perfusion, underscoring its therapeutic capacity to resolve the no-reflow phenomenon.

**Figure 5.**
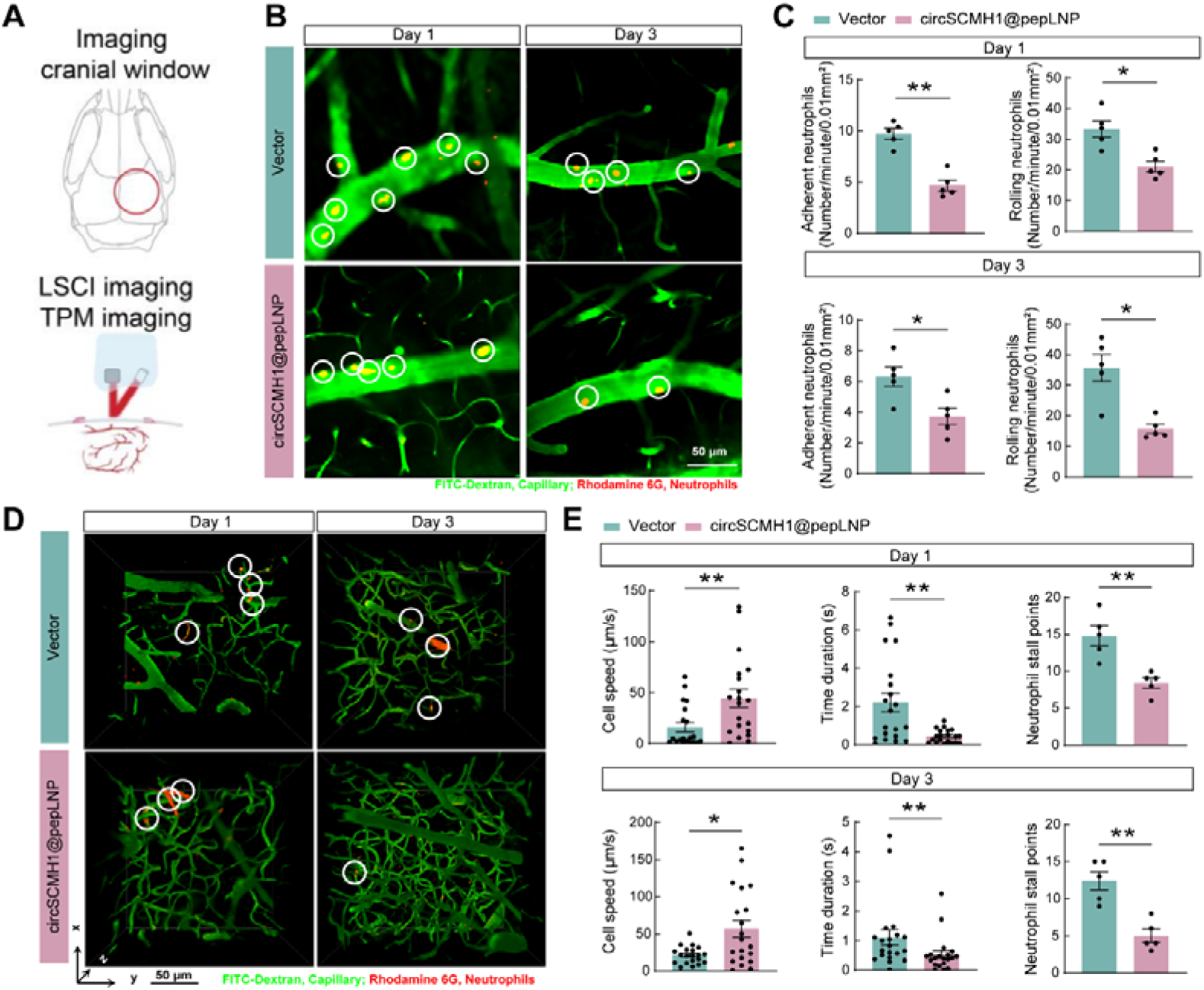
Intravital two-photon imaging reveals that circSCMH1@pepLNP mitigates neutrophil adhesion and alleviates capillary stalling *in vivo*. (A) Schematic illustration of the cranial window implantation site and the experimental setup for intravital two-photon microscopy (TPM) and laser speckle contrast imaging (LSCI). **(B-C)** Representative intravital two-photon images (B) and corresponding quantitative analysis (C) of neutrophil adhesion and rolling dynamics within the cortical microvasculature at days 1 and 3 post-reperfusion. The vascular lumen was perfused with FITC-Dextran (green), and neutrophils were labeled with Rhodamine 6G (red/yellow due to colocalization). White circles highlight representative adherent or rolling neutrophils. Scale bar: 50 μm. N = 5 mice per group (data averaged from 5 FOVs per mouse). \**p*<0.05 and \*\**p*<0.01 using Student’s t test. **(D-E)** Representative 3D reconstructed vascular networks derived from two-photon z-stacks (230 × 230 × 150 μm³) (D) and quantitative tracing analysis of neutrophil dynamics (E) at days 1 and 3 post-reperfusion. White circles in (D) indicate stalled neutrophils plugging the capillary segments. Quantitative parameters in (E) include individual neutrophil migration speed, duration of capillary stalling (time duration), and the cumulative count of neutrophil stall points within a defined field of view (FOV, 230 × 230 μm²). N = 20 individual cell traces from 5 mice per group for cell speed and duration; n = 5 mice per group for neutrophil stall points. \**p*<0.05 and \*\**p*<0.01 using Student’s t test. FOV, field of view. LSCI, laser speckle cortical imaging.

## Discussion

The clinical efficacy of EVT in large vessel occlusion acute ischemic stroke is frequently compromised by FR, a phenomenon driven by post-reperfusion microvascular thromboinflammation and capillary stalling. In this study, we bridge this translational gap by elucidating the pathogenic relevance of circSCMH1 downregulation in clinical FR and employing a targeted nanomedicine approach to spatiotemporally unlock its acute therapeutic potential. Clinical analysis revealed circSCMH1 levels are significantly decreased in both the thrombi and plasma of LVO-AIS patients with FR, and that this systemic reduction correlates negatively with the circulating NET burden. Furthermore, we engineered a pepLNP system to specifically deliver circSCMH1 to neutrophils. This targeted intervention exerted potent anti-thromboinflammatory effects by abolishing NET formation, alleviating microvascular obstruction, and restoring cerebral perfusion. Importantly, our work identifies a previously unrecognized anti-thromboinflammatory function of circSCMH1, thereby conceptualizing a dual-phase therapeutic paradigm that links hyperacute thromboinflammation suppression to long-term neurovascular repair.

While EVT has revolutionized the recanalization of large cerebral arteries, recent clinical evidence underscores that macroscopic recanalization alone is frequently insufficient to achieve meaningful tissue-level reperfusion. This persistent microvascular failure has spurred the exploration of novel clinical paradigms, such as EVT combined with adjunctive intra-arterial thrombolysis or targeted anti-thrombotic therapies, aimed at rescuing the ischemic microcirculation.^17–20^ However, focusing solely on residual clot dissolution overlooks the profound, neutrophil-driven thromboinflammation triggered immediately upon reperfusion.^21^ Consequently, the contemporary theoretical framework for stroke management is shifting toward a more comprehensive, dual-phase intervention strategy: one that must actively neutralize the hyperacute thromboinflammatory cascade while simultaneously fostering delayed neurovascular regeneration.^22^ For decades, the failure to translate preclinical neuroprotective agents into clinical practice has been largely attributed to their inability to navigate this spatiotemporal complexity.^8,23^ Most traditional therapies singularly target either early injury mechanisms or late-stage repair within a rigid, narrow therapeutic window, inevitably leading to suboptimal global efficacy.^24,25^ The targeted nanomedicine strategy proposed in our study is strategically designed to potentially navigate this translational bottleneck. By simultaneously engaging the suppression of hyperacute thromboinflammation via NETosis inhibition and the preservation of long-term neurovascular recovery, circSCMH1@pepLNP offers a promising framework to cover the entire pathophysiological spectrum of ischemic stroke.

Current clinical investigations into adjunctive therapies post-EVT predominantly focus on intra-arterial thrombolytics, such as alteplase or tenecteplase, and antiplatelet agents like tirofiban.^19,26^ These pharmacological strategies primarily target residual fibrin networks and platelet aggregation to maintain large-vessel patency. While these approaches have shown promise in specific clinical scenarios, the complex pathophysiology of microvascular no-reflow extends beyond simple thrombus dissolution.^27,28^ Our study rationalizes targeting neutrophils as a critical complementary strategy, given their distinct dual pathogenic role in the hyperacute phase: functioning simultaneously as mechanical obstructers of the capillary bed and primary drivers of thromboinflammation via rapid NETosis.^21,29^ However, systemic administration of non-targeted nanotherapeutics typically results in extensive clearance by the reticuloendothelial system, limiting their bioavailability at the ischemic core. To overcome this fundamental limitation, we employed the FPR-targeted pepLNP system to actively intercept circulating neutrophils prior to their microvascular infiltration. Our intravital imaging data provide direct visual confirmation of this targeted efficacy, demonstrating that pepLNP-mediated intervention significantly reduces early neutrophil adhesion and subsequent capillary stalling. This specific cellular modulation effectively resolves the dual pathogenic threat of neutrophils, thereby preserving cortical microvascular perfusion and overcoming the delivery constraints of conventional neuroprotectants.

Beyond the strategic advantages of targeted delivery, our study significantly expands the functional landscape of circSCMH1, positioning it as a pivotal molecular bridge within the biphasic pathology of ischemic stroke. Building upon our laboratory’s extensive prior work establishing circSCMH1 as a key driver of late-stage neuroplasticity, angiogenesis, and astrocytic restoration, we now uncover an unexpected dimension to its biology in the hyperacute phase.^14,30–32^ While our earlier studies focused on its regenerative capacity, the current findings reveal that circSCMH1 also functions as a potent, endogenous negative regulator of innate immune activation. Importantly, the targeted intracellular delivery via pepLNP was essential to unlock this acute anti-thromboinflammatory potential, allowing circSCMH1 to exert its protective effects in a time window previously thought inaccessible. This synergistic combination of molecular pleiotropy and precise delivery directly addresses the challenge of spatiotemporal complexity in stroke therapy.^33,34^ By demonstrating that neutrophil-targeted circSCMH1 effectively suppresses early NETosis while retaining its inherent reparative properties, we identify a unified strategy that addresses the full continuum of ischemic damage. These findings provide a strong mechanistic foundation for a "single-agent, dual-phase" therapeutic paradigm, effectively bridging the critical gap between acute thromboinflammatory mitigation and sustained neurological recovery.

Despite these promising findings, several limitations warrant consideration. Primarily, the precise molecular mechanism by which circSCMH1 inhibits NETosis remains to be fully elucidated. While our previous work established that circSCMH1 promotes vascular repair via FTO-dependent m6A demethylation, we did not investigate whether this specific epigenetic pathway contributes to its acute anti-NETotic effects. Future studies should prioritize identifying the downstream molecular targets of circSCMH1 within neutrophils, potentially involving metabolic reprogramming or signaling modulation, to clarify its exact mode of action. Furthermore, the efficacy of the circSCMH1@pepLNP delivery system was evaluated exclusively in murine models, necessitating extensive further investigation into its clinical translatability. Key future directions include optimizing the nanoparticle formulation for human scaling, rigorously assessing its pharmacokinetic and pharmacodynamic profiles, and evaluating potential systemic toxicities, which represent critical hurdles that must be overcome to advance nanomedicines toward clinical application.

Our study identifies circSCMH1 downregulation as a potential driver of FR in LVO-AIS. By employing a targeted delivery strategy, we successfully uncovered a potent early anti-thromboinflammatory function of this transcript, providing a critical mechanistic complement to its established late-stage neuroprotective effects. These findings expand our understanding of circSCMH1 as a dual-phase modulator and support a single-formulation therapeutic strategy for comprehensive stroke management. Ultimately, this work proposes a novel adjunctive approach for EVT designed to mitigate microvascular reperfusion injury and improve clinical outcomes.

## Author contributions

L.X., X.H. and S.L. recruited AIS patients, collected clinical patient blood samples, isolated neutrophils. L.X. and Z.Z. performed animal experiments, morphological experiments and statistical analysis. Y.J. and Y.B. prepared the circSCMH1@pepLNP.

B.H. contributed to the clinical sample analysis. Last, H.Y., B.H, and X.H., conceived the project, organized the study, and supervised the experiments.

## Funding support

This work was supported by grants from Noncommunicable Chronic Diseases-National Science and Technology Major Project (2024ZD0526600/2024ZD0526601), the National Natural Science Foundation of Distinguished Young Scholars (82025033), the National Natural Science Foundation of China (82230115, 82404602, 82572433, 82504772), China Postdoctoral Science Foundation (2024M760452), Research Personnel Cultivation Programme of Zhongda Hospital Southeast University (CZXM-GSP-RC85), Zhongda Hospital Affiliated to Southeast University, Jiangsu Province High-Level Hospital Construction Funds (GSP-LCYJFH28).

## Competing interests

The authors declare no competing interests.

## Notes

### Competing Interest Statement

The authors have declared no competing interest.

