## Supplemental_figures for "Neutrophil-targeted delivery of circSCMH1 unlocks an acute anti-thromboinflammatory function to restore microvascular perfusion in stroke": Supplemental Figures.pdf

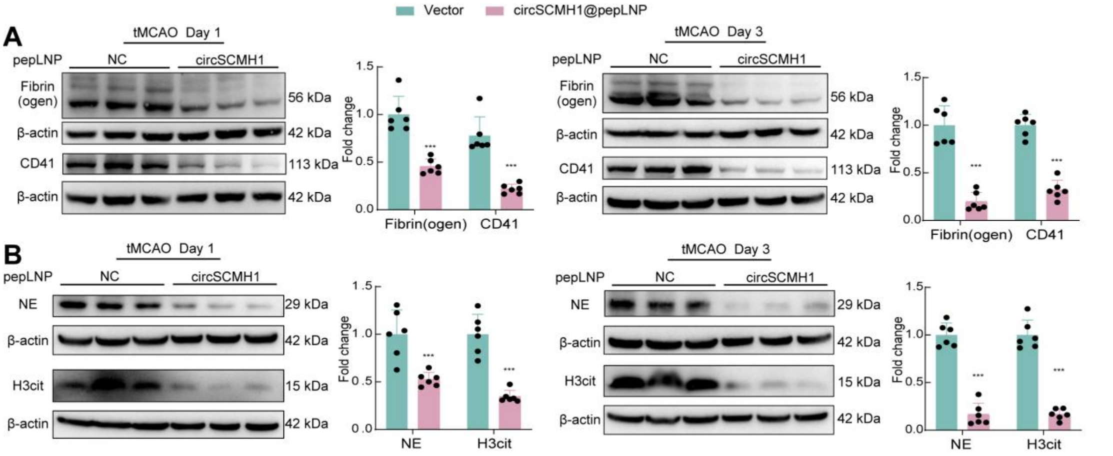


**Supplemental Figure 1. Western blot analysis of ischemic brain from tMCAO mice at days 1 and 3 post-reperfusion.** (**A**)Western blot analysis of Fibrin(ogen) and CD41 expression in the ischemic brain from tMCAO mice at days 1 and 3 post-reperfusion. n= 6 mice in each group. ****p*<0.001 using Student's t test. (**B**)Western blot analysis of NE and H3cit expression in the ischemic brain from tMCAO mice at days 1 and 3 post-reperfusion. n= 6 mice in each group. ****p*<0.001 using Student's t test.

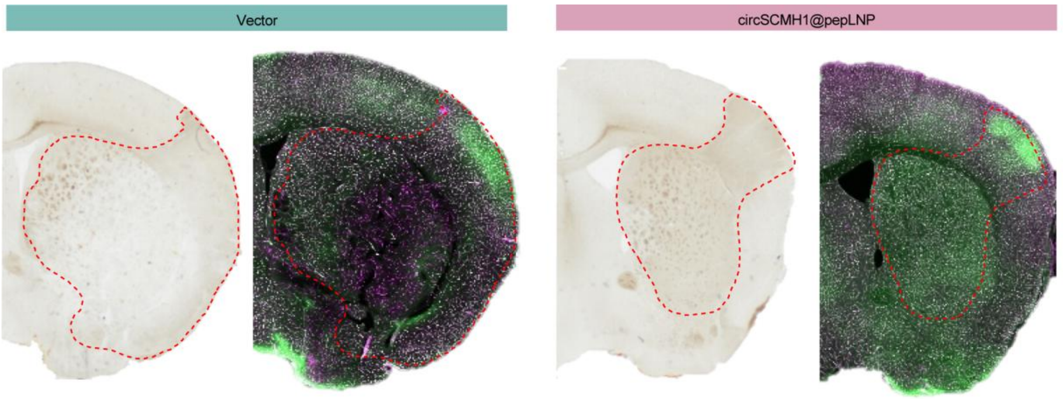


**Supplemental Figure 2. Representative scanning immunofluorescence images of the ischemic brain from tMCAO mice.** Non-reflow microvessels only labeled by red LEL, reflow microvessels both labeled red and green LEL. LEL: *lycopersicon esculentum* (tomato) lectin.
